# Structural Design of Chimeric Antigens for Multivalent Protein Vaccines

**DOI:** 10.1101/187336

**Authors:** Sarah Hollingshead, Ilse Jongerius, Rachel M. Exley, Steven Johnson, Susan M. Lea, Christoph M. Tang

## Abstract

The development of prophylactic vaccines against pathogenic bacteria is a major objective of the World Health Organisation. However, vaccine development is often hindered by antigenic diversity and the difficulties encountered manufacturing immunogenic membrane proteins. Here, we employed structure-based design as a strategy to develop <u>Ch</u>imeric <u>A</u>ntigens (ChAs) for subunit vaccines. ChAs were generated against serogroup B *Neisseria meningitidis* (MenB), the predominant cause of meningococcal disease in the Western hemisphere. MenB ChAs exploit the lipoprotein factor H binding protein (fHbp) as a molecular scaffold to display the immunogenic VR2 epitope from the integral membrane protein PorA. Structural analyses demonstrate fHbp is correctly folded and that PorA VR2 epitope adopts an immunogenic conformation. In mice, ChAs elicit antibodies directed against fHbp and PorA, with antibody responses correlating to protection against meningococcal disease. ChAs offer a novel approach for generating multivalent subunit vaccines, containing of epitopes from integral membrane proteins, whose composition can be selected to circumvent pathogen diversity.

## Introduction

Toxoid and capsule-based vaccines have saved millions of lives caused by bacterial pathogens (Andre, Booy et al., 2008). For example, toxoid based vaccines have virtually eliminated diphtheria and tetanus in wealthy countries (Arístegui, Usonis et al., 2003, Schmitt, von Kries et al., 2001), while capsule-based vaccines have significantly reduced disease caused by *Haemophilus influenza* (Murphy, 2015), *Streptococcus pneumonia* (Keller, Robinson et al., 2016), and certain strains of *Neisseria meningitidis* (Maiden, 2013). However, significant challenges remain in developing vaccines against pathogens for which toxoid and capsule-based vaccines are not viable. These pathogens include non-typeable strains of *H. influenza* and *S. pneumonia* (Keller et al., 2016, Murphy, 2015), unencapsuated pathogens such as *Neisseria gonorrhoeae* and *Moraella catarrhalis* (HPA, 2013, HPA, 2016, Jerse, Bash et al., 2014, Schaller, Troller et al., 2006) and encapsulated serogroup B *N. meningitidis*, for which a capsule based vaccine is not feasible (Finne, Leinonen et al., 1983). Given the exponential rise in the emergence of multi-drug resistant bacteria (WHO, 2014, Wi, Lahra et al., 2017), new approaches for vaccine development are paramount. However, strategies for generating successful vaccines are hampered by pathogen diversity (Telford, 2008) and the difficulties associated with presenting epitopes from membrane-embedded surface proteins to the immune system (Carpenter, Beis et al., 2008).

Two main approaches have been used to develop vaccines against serogroup B *N. meningitidis*; outer membrane vesicle vaccines (OMVVs) and recombinant protein subunit vaccines (RPSVs). OMVVs were first developed around thirty years ago (Bjune, Høiby et al., 1991, O’Hallahan, Lennon et al., 2004, Sierra, Campa et al., 1991). The immunodominant antigen in meningococcal OMVVs is PorA (Saukkonen, Leinonen et al., 1989), an abundant outer-membrane porin with eight surface-exposed loops (Blake & Gotschilch, 1987, Judd, 1989). Loops one and four are termed Variable Region 1 and 2 (VR1 and VR2), respectively, as they elicit immune responses and are subject to antigenic variation (McGuinness, Barlow et al., 1990, van der Ley, Heckels et al., 1991). The VR2 loop dominates PorA-specific immunity elicited by OMVVs, which offer limited or no cross-protection against strains expressing PorA with a different VR2 (Holst, Martin et al., 2009, Martin, Ruijne et al., 2006, Michaelsen, Ihle et al., 2003). To broaden coverage, OMVVs have been developed containing multiple PorAs (Claassen, Meylis et al., 1996, van den Dobbelsteen, van Dijken et al., 2004, van der Ley, van der Biezen et al., 1995), selected for their prevalence in circulating strains (Luijkx, van Dijken et al., 2003, van den Dobbelsteen et al., 2004). However, OMVVs present complex manufacturing and regulatory issues (Ulmer, Valley et al., 2006). More recently, RPSVs Bexsero and Trumenba were developed. These contain the key meningococcal antigen factor H binding protein (fHbp), a lipoprotein composed of two β-barrels that tightly bind domains 6 and 7 of human complement Factor H (CFH) (Madico, Welsch et al., 2006, Pizza, Scarlato et al., 2000, Schneider, Exley et al., 2006, Schneider, Prosser et al., 2009). fHbp is antigenically variable; public databases contain over 900 different fHbp variants (Jolley & Maiden, Jolley & Maiden, 2010), which fall into three variant groups or two subfamilies: V1 (subfamily B), V2 and V3 (both subfamily A) (Brehony, Wilson et al., 2009, Murphy, Andrew et al., 2009). In general, immunisation with a particular fHbp induces cross-protection against strains expressing fHbp belonging to the same, but not a different, variant group; although there is significant cross-protection between variant groups 2 and 3 (subfamily A) fHbps (Fletcher, Bernfield et al., 2004, Masignani, Comanducci et al., 2003). Bexsero contains a single fHbp variant (V1.1), with two other recombinant antigens as well as an OMV (Serruto, Bottomley et al., 2012), whilst Trumenba is solely composed of two fHbp variants (V1.55 and V3.45) (Green, Eiden et al., 2016). Antigens in Bexsero and Trumenba have exact sequence matches to only 36% and 4.8% of serogroup B *N. meningitidis* disease isolates currently circulating in the UK, respectively (Brehony, Hill et al., 2015, Jolley & Maiden), leading to concerns about their ability to provide broad coverage against an antigenically diverse pathogen.

We employed a structure-based approach to generate <u>Ch</u>imeric <u>A</u>ntigen<u>s</u> (ChAs) against serogroup B *N. meningitidis*. ChAs exploit fHbp as a molecular scaffold to present the surface exposed PorA VR2 loop, which is achieved by inserting the VR2 loop into a β-turn region in fHbp. ChAs retain epitopes from both fHbp and PorA, and can elicit functional immune responses against both antigens. We demonstrate integration of a VR2 loop does not alter the overall architecture of fHbp and that the VR2 loop folds into a conformation recognized by a bactericidal mAb. We provide proof-in-principle that ChAs can be used to display selected epitopes from integral membrane proteins, such as PorA. ChAs incorporate epitopes from multiple antigens into a single vaccine antigen, which can be selected to circumvent pathogen antigenic diversity. Furthermore, ChAs contain epitopes from integral membrane proteins, which have previously hindered vaccine development, owing to the difficulties encountered during manufacture.

## Results

### Design and construction of chimeric fHbp:PorAs

Immunisation with *N. meningitidis* proteins fHbp and PorA elicits bactericidal antibody responses, which provide a correlate of protection against meningococcal disease (Green et al., 2016, Serruto et al., 2012) (**Figure 1A**). fHbp is a lipoprotein that expresses as a soluble protein in *Escherichia coli* following removal of the N-terminal lipobox motif (Masignani et al., 2003). As an extracellular loop, the PorA VR2 is likely soluble when expressed separately from the integral membrane regions of PorA. We exploited soluble fHbp variant 1.1 (V1.1) as a molecular scaffold to display the PorA VR2 loop, P1.16. The PorA VR2 loop P1.16 (YYTKDTNNNLTLV) was inserted into six different β-turn regions in fHbp. At each site the PorA VR2 loop was inserted, a single amino acid was deleted from the fHbp scaffold (**Figure 1B**). The resulting fHbp:PorA ChAs were named according to a scheme in which fHbp^V1.1^:PorA^151/P1.16^ denotes fHbp V1.1 with PorA VR2 loop P1.16 inserted at residue 151 of fHbp (**Table S1**). The ChAs all express to high levels in *E. coli* and were purified by nickel affinity chromatography. Western blot analyses confirm all ChAs retain epitopes recognised by an α-P1.16 mAb and α-fHbp pAbs (**Figure 1C**).

**Figure 1.**
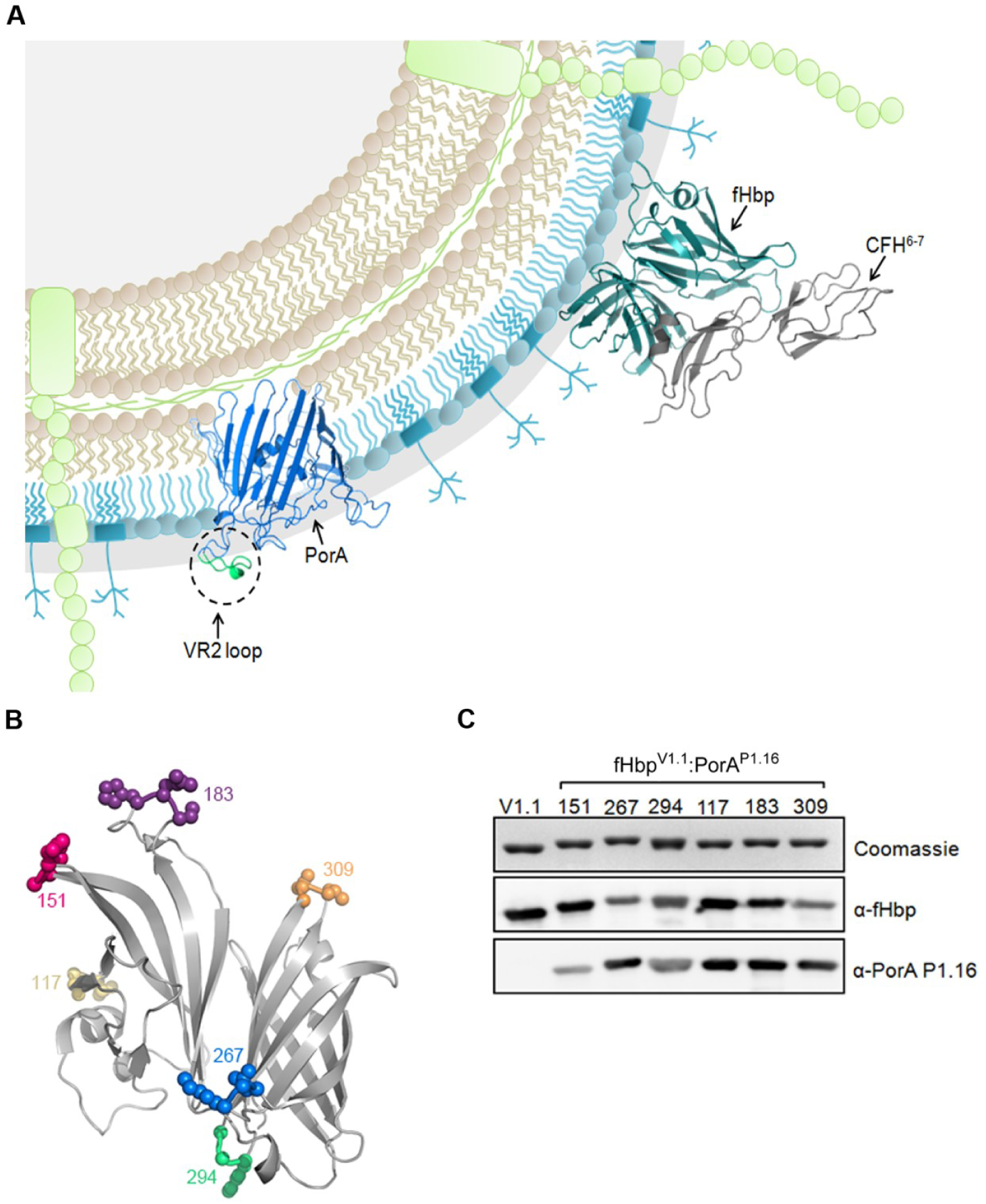
Structure based design of fHbp:PorA ChAs. (**A**) Schematic of meningococcal cell surface, depicting the key surface exposed antigens fHbp and PorA. (**B**) Location of fHbp residues replaced with the PorA P1.16 VR2 loop. (**C**) Analysis of recombinant ChAs by SDS-PAGE and Western blot. Immunoblots are probed with α-V1.1 fHbp pAb and α-PorA P1.16 mAb.

### fHbp:PorAs are stable and can bind CFH

Stability of an antigen is an important property of a vaccine, and insertion of PorA epitopes might disrupt the overall structure of the ChA scaffold. Therefore, we determined the thermal stability of ChAs by differential scanning calorimetry (DSC, **Table 1**). Insertion of a PorA loop into the N-or C-terminal β-barrel of fHbp decreased the thermal stability of that β-barrel by 1.0°C to 15.5°C, with little or no effect on the other β-barrel. Overall, the lowest measured melting temperature (T_m_) of any β-barrel was 60.5°C, which is considerably higher than the N-terminal T_m_ of V3.45 (41°C), one of the fHbps in Trumenba^®^ (**Table 1**). A key property of fHbp is its ability to bind CFH (Madico et al., 2006, Schneider et al., 2006, Schneider et al., 2009) (**Figure S1A**). Therefore, surface plasmon resonance (SPR) was used to determine the affinity of each ChA for domains 6 and 7 of CFH (**Table 1**). Most ChAs bind CFH at high affinity, indicating the fHbp scaffold retains its function. The exceptions were fHbp^V1.1^:PorA^183/P1.16^, to which there was no detectable CFH binding, and fHbp^V1.1^:PorA^267/P1.16^, to which CFH binding was reduced by approximately eight fold. In these two ChAs, the VR2 loop is situated in the region of fHbp that engages CFH (**Figure S1B**), potentially inhibiting CFH binding.

**Table 1:** Stability of fHbp:PorA ChAs and affinity for CFH

| Protein | Cp (kcal mole<sup>-1</sup> °C<sup>-1</sup>) |  | Kd (nM) |
| --- | --- | --- | --- |
|  | N-terminal T<sub>m</sub> | C-Terminal T<sub>m</sub> |  |
| fHbp V1.1 | 69.8 | 87.9 | 3.6 |
| fHbp<sup>V1.1</sup>:PorA<sup>151/P1.16</sup> | 60.5 | 87.3 | 2.3 |
| fHbp<sup>V1.1</sup>:PorA<sup>267/P1.16</sup> | 68.4 | 75.7 | 28 |
| fHbp<sup>V1.1</sup>:PorA<sup>294/P1.16</sup> | 61.7 | 72.4 | 2.7 |
| fHbp<sup>V1.1</sup>:PorA<sup>117/P1.16</sup> | 65.0 | 87.9 | 2.3 |
| fHbp<sup>V1.1</sup>:PorA<sup>183/P1.16</sup> | 68.8 | 88.0 | NB |
| fHbp<sup>V1.1</sup>:PorA<sup>309/P1.16</sup> | 69.0 | 76.9 | 5.5 |
| fHbp V3.45 | 41.0 | 83.0 | - |
| fHbp<sup>V1.4</sup>:PorA<sup>151/P1.1.10_1</sup> | 54.0 | 89.0 | - |
| fHbp<sup>V1.4</sup>:PorA<sup>151/P1.14</sup> | 55.0 | 88.0 | - |
| fHbp<sup>V1.4</sup>:PorA<sup>151/P1.15</sup> | 55.0 | 89.0 | - |
| fHbp<sup>V3.45</sup>:PorA<sup>158/P1.4</sup> | 40.0 | 81.0 | - |
| fHbp<sup>V3.45</sup>:PorA<sup>158/P1.9</sup> | 39.0 | 80.0 | - |

Melting temperature, T<sub>m</sub>; Kd, dissociation constant; non binding, NB.

### fHbp:PorA ChAs elicit protective immune responses

To examine the ability of ChAs to elicit immune responses, groups of CD1 mice were immunized with ChAs using alum or monophospholipid A (MPLA) as the adjuvant (**Figure 2A**). Post immunisation, sera were obtained from mice and pooled. Immune responses were assessed against *N. meningitidis* H44/76, a serogroup B strain which expresses V1.1 fHbp and PorA VR2 P1.16. We constructed isogenic strains to define immune responses directed against fHbp (H44/76Δ*porA*) and PorA (H44/76Δ*fHbp*), as well as a H44/76Δ*fHbp*Δ*porA* control. Western blot analyses of lysates from these strains demonstrate all ChAs elicit antibodies that recognise both fHbp and PorA (**Figure 2B and 2D**), except for fHbp^V1.1^:PorA^267/P1.16^/MPLA which generates sera that only recognises fHbp (**Figure 2D**).

**Figure 2.**
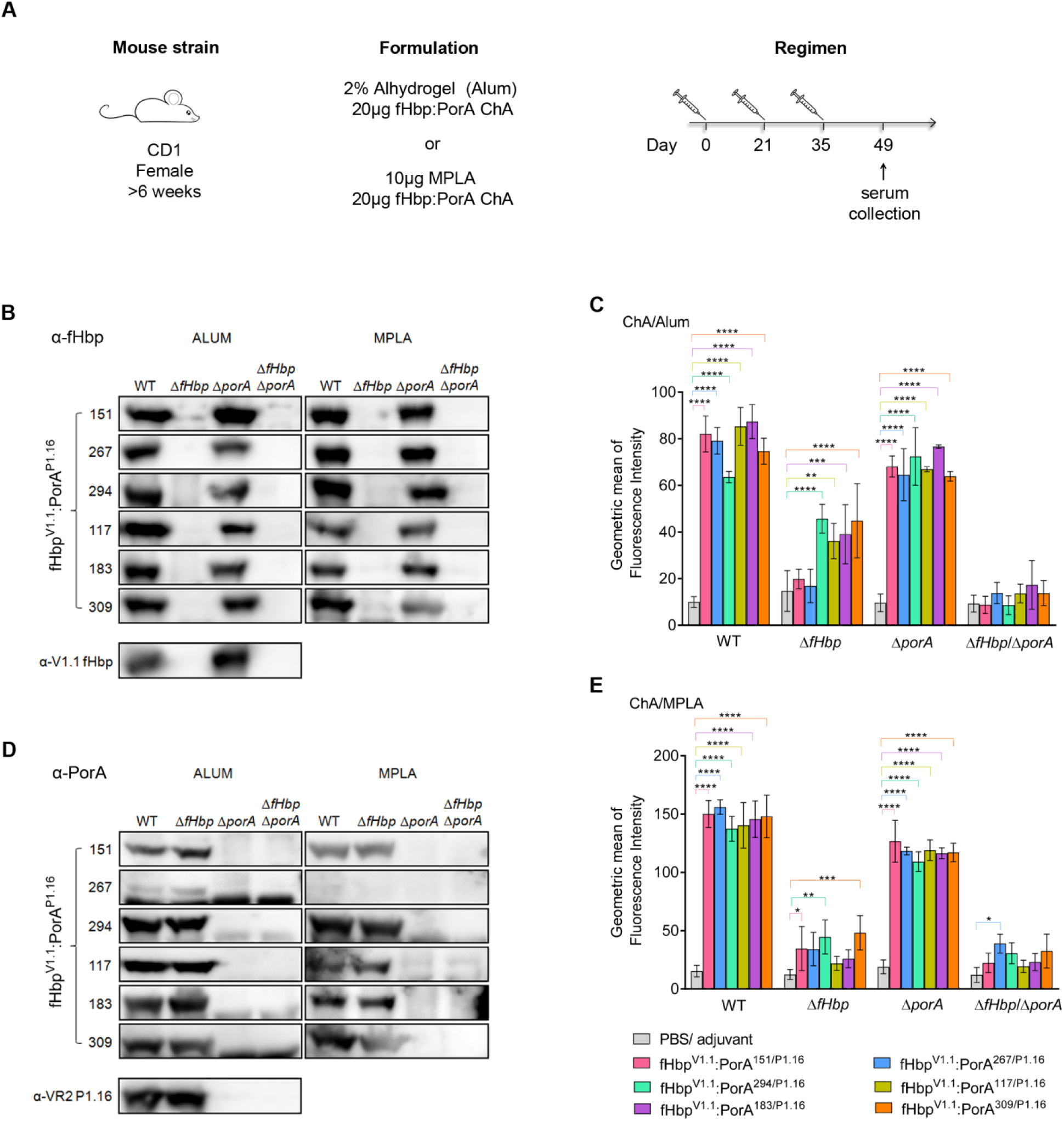
Detection of fHbp and PorA by fHbp:PorA ChA antisera. (**A**) Immunisation strategy. Mice (*n*=8 per group) were subcutaneously immunized three times with each combination of ChA/Adjuvant. (**B** and **D**) Western blots of *N. meningitidis* whole cell lysates probed with each set of ChA/adjuvant antisera; (**B**) Detection of fHbp, (**D**) Detection of PorA. (**C** and **E**) Flow cytometry analysis showing binding of each set of ChA/adjuvant antisera to *N. meningitidis* H44/76 strains WT, Δ*fHbp*, Δ*porA* and Δ*fHbp*Δ*porA*. SD of independent assays (*n*=3) is indicated. Two-way ANOVA and Dunnett’s method of multiple comparison were used to compare the fluorescence intensity of ChA antisera with PBS control sera (**C** and **E**, * *p*≤0.05, ** *p*≤0.01, *** *p*≤0.001, **** *p*≤0.0001).

Next, we used flow cytometry to assess the ability of antibodies raised during ChA immunisations to recognise fHbp and PorA on the surface of *N. meningitidis.* We detected antibody specific binding to fHbp and PorA by flow cytometry, using *N. meningitidis* strains H44/76Δ*porA* and H44/76Δ*fHbp*, respectively. Binding was determined by comparison with control sera, obtained from mice immunised with PBS and adjuvant alone. Antisera raised against each ChA detected fHbp on the bacterial surface (*p*≤0.0001, **Figure 2C and 2E**), and certain antisera also demonstrated significant binding to PorA: antisera from mice immunised with alum and fHbp^V1.1^:PorA^294/P1.16^, fHbp^V1.1^:PorA^117/P1.16^, fHbp^V1.1^:PorA^183/P1.16^ or fHbp^V1.1^:PorA^309/P1.16^ (*p*≤0.01, **Figure 2C**), and from mice immunised with MPLA and fHbp^V1.1^:PorA^151/P1.16^, fHbp^V1.1^:PorA^294/P1.16^ or fHbp^V1.1^:PorA^309/P1.16^ (**Figure 2E**, *p*≤0.05). Significant binding (*p*≤0.05), is observed with fHbp^V1.1^:PorA^267/P1.16^/MPLA antisera to the H44/76Δ*fHbp*Δ*porA* negative control strain, which is due to non-specific binding (**Figure S2**).

The serum bactericidal assay (SBA) assesses the ability of antibodies to initiate complement-mediated lysis of *N. meningitidis.* When using baby rabbit complement, an SBA titre of ≥8 is an accepted correlate of protective immunity against *N. meningitidis* (Andrews, Borrow et al., 2003). SBAs conducted with each set of pooled ChA/adjuvant antisera and wild-type *N. meningitidis* H44/76 all had titres of ≥128 (**Figure 3A**). Significantly higher titres (*p*≤0.05) were observed for antisera raised against fHbp^V1.1^:PorA^151/P1.16^, fHbp^V1.1^:PorA^117/P1.16^, fHbp^V1.1^:PorA^183/P1.16^ and fHbp^V1.1^:PorA^309/P1.16^ when MPLA, rather than alum, was used as the adjuvant.

**Figure 3.**
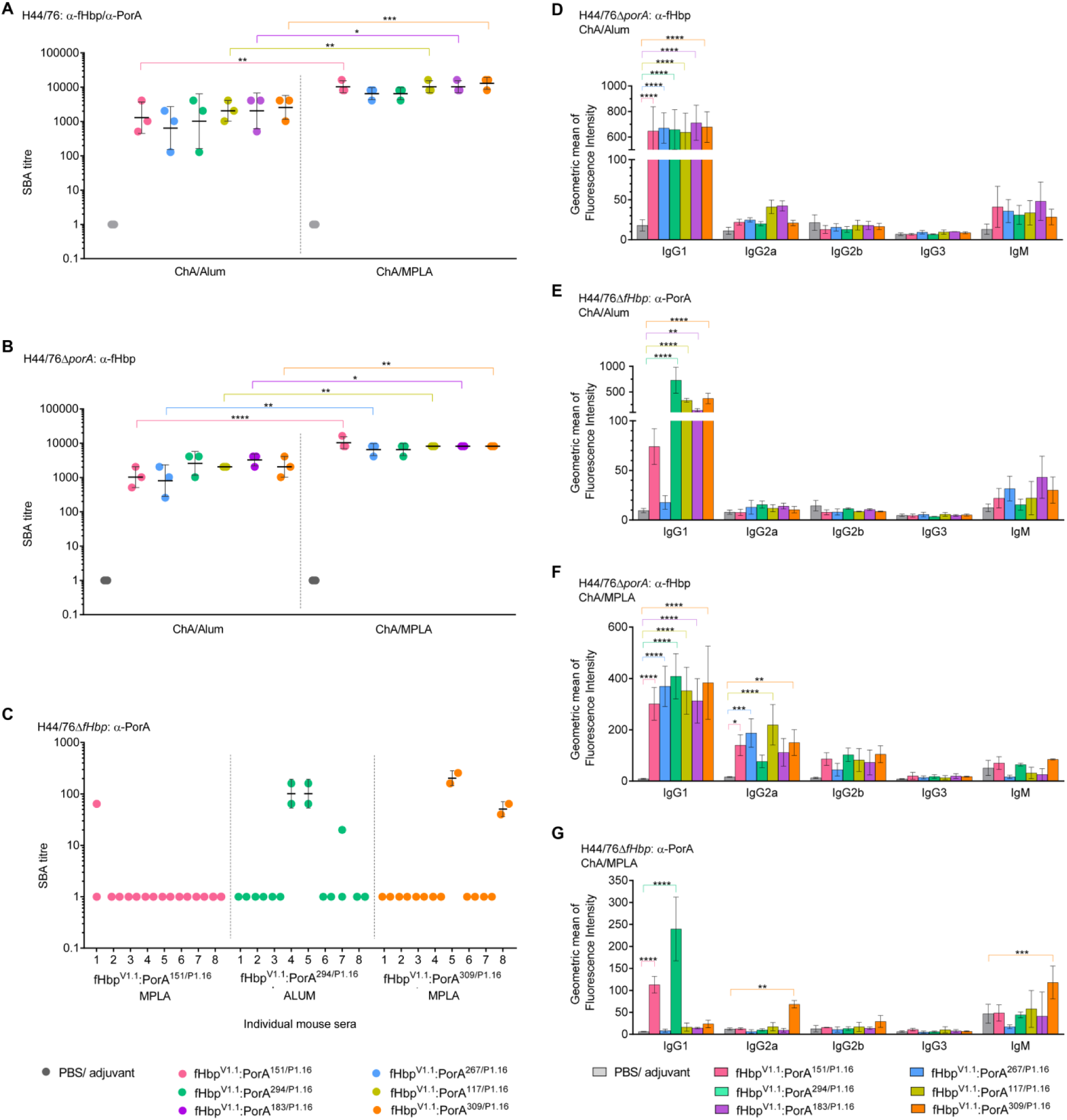
Immunogenicity of fHb:PorA ChAs. Titres from SBAs performed with *N. meningitidis* strains H44/76. (**A**, *n*=3), H44/76Δ*porA* (**B**, *n*=3), and H44/76Δ*fHbp* (**C**, *n*=2) and ChA/adjuvant antisera. SBAs with H44/76 and H44/76Δ*porA* were conducted using pooled ChA/adjuvant antisera, and SBAs with H44/76Δ*fHbp* were conducted with ChA/adjuvant antisera from individual mice. Geometric mean and SD of independent assays (*n*>2) are indicated. Flow cytometry was used to detect binding of mouse isotypes IgG1, IgG2a, IgG2b, IgG3 and IgM in ChA antisera to *N. meningitidis* strains H44/76Δ*porA* (**D** and **F**) and H44/76Δ*fHbp* (**E** and **G**). Mouse isotypes binding to *N. meningitidis* were detected with isotype specific secondary antibodies. SD of independent assays (*n*=3) is indicated. Two-way ANOVA and Dunnett’s method of multiple comparison were used to compare SBA titres from pooled antisera (**A** and **B**) and ChA antisera to PBS control sera (**D**-**G**) (* *p*≤0.05, ** *p*≤0.01, *** *p*≤0.001, **** *p*≤0.0001).

To test for fHbp-specific responses, we performed SBAs with *N. meningitidis* H44/76Δ*porA*; immunisations with every ChA/adjuvant generated significant SBA titres (**Figure 3B**). To evaluate α-PorA responses, we initially performed SBAs with pooled antisera. However, PorA-dependent complement mediated lysis was not detected with pooled antisera. Therefore, we examined PorA-dependent responses in individual mice. SBA titres were detected with antisera raised against fHbp^V1.1^:PorA^294/P1.16^ when alum was used as the adjuvant, and with antisera raised against fHbp^V1.1^:PorA^151/P1.16^ (a low level of SBA from a single mouse), or fHbp^V1.1^:PorA^309/P1.16^ when MPLA was the adjuvant (**Figure 3C**). Although PorA was detected on the surface of *N. meningitidis* by antisera from mice immunised with alum and fHbp^V1.1^:PorA^117/P1.16^ or fHbp^V1.1^:PorA^183/P1.16^, or with MPLA and fHbp^V1.1^:PorA^294/P1.16^ (**Figure 2C and 2E**), these antisera did not have PorA-dependent SBA titres.

To activate the classical pathway, bound immunoglobulin (Ig) must recruit the C1q subunit of C1 (Frank, Joiner et al., 1987). The ability of Ig classes to bind C1q varies; a single IgM can be sufficient for C1q recruitment (Poon, Phillips et al., 1985), while several IgGs must be bound in close proximity and in a particular conformation(Burton, 1990, Hughes-Jones & Gardner, 1979, Sledge & Bing, 1973). Therefore, we examined which Ig isotypes are elicited by ChAs. Flow cytometry demonstrates that IgG1 is the main Ig in immune sera that binds the surface of *N. meningitidis* (**Figure 3D-3G**). When compared with sera from mice immunised with PBS/adjuvant alone, all ChAs elicit significant α-fHbp IgG1 responses (**Figure 3D and 3F**, *p*≤0.0001). However, significant IgG1 binding to PorA on the surface of *H44/76*Δ*fHbp* was observed only with antisera raised against fHbp^V1.1^:PorA^294/P1.16^, fHbp^V1.1^:PorA^117/P1.16^, fHbp^V1.1^:PorA^183/P1.16^ or fHbp^V1.1^:PorA^309/P1.16^ with alum (*p*≤0.01, **Figure 3E**), and fHbp^V1.1^:PorA^151/P1.16^ or fHbp^V1.1^:PorA^294/P1.16^ with MPLA (*p*≤0.0001, **Figure 3G**). Interestingly, there was no detectable IgG1 binding to PorA using sera raised against fHbp^V1.1^:PorA^309/P1.16^/MPLA, against which two mice had α-PorA SBA titres (**Figure 3C**); instead the sera contained significant levels of α-PorA IgG2a and IgM (*p*≤0.01, **Figure 3G**).

### ChAs retain the architecture of the fHbp scaffold and PorA loop

To further characterise the fHbp:PorA ChAs, we determined the atomic structures of the P1.16 VR2 loop in residues 151, 294, and 309 of a V1 fHbp, all of which generated SBA titres. The structures were solved using molecular replacement to resolutions of 2.9 Å for fHbp^V1.4^:PorA^151/P1.16^, 3.7 Å for fHbp^V1.1^:PorA^294/P1.16^ and 2.6 Å for fHbp^V1.4^:PorA^309/P1.16^. Alignment of all fHbp scaffolds with V1.1 fHbp (**Figure S1C**) showed good agreement (RMSDs range between 0.447 and 0.614), demonstrating that the fHbp scaffold is not perturbed by insertion of the VR2 loop. In the ChAs, the VR2 loops adopt a β-turn conformation that extends away from the main body of fHbp, without interacting with the fHbp scaffold (**Figure 4A**). Of note, the conformations of PorA VR2 P1.16 in the ChAs (fHbp^V1.4^:PorA^151/P1.16^, fHbp^V1.1^:PorA^294/P1.16^, fHbp^V1.4^:PorA^309/P1.16^) and in a complex with the Fab fragment (van den Elsen, Herron et al., 1997) all align with good agreement (**Figure 4B**, RMSDs range between 0.346 - 0.970), thus demonstrating that the PorA VR2 P1.16 loops in the ChAs adopt a conformation that can induce bactericidal antibody responses (Oomen, Hoogerhout et al., 2003, Oomen, Hoogerhout et al., 2005).

**Figure 4.**
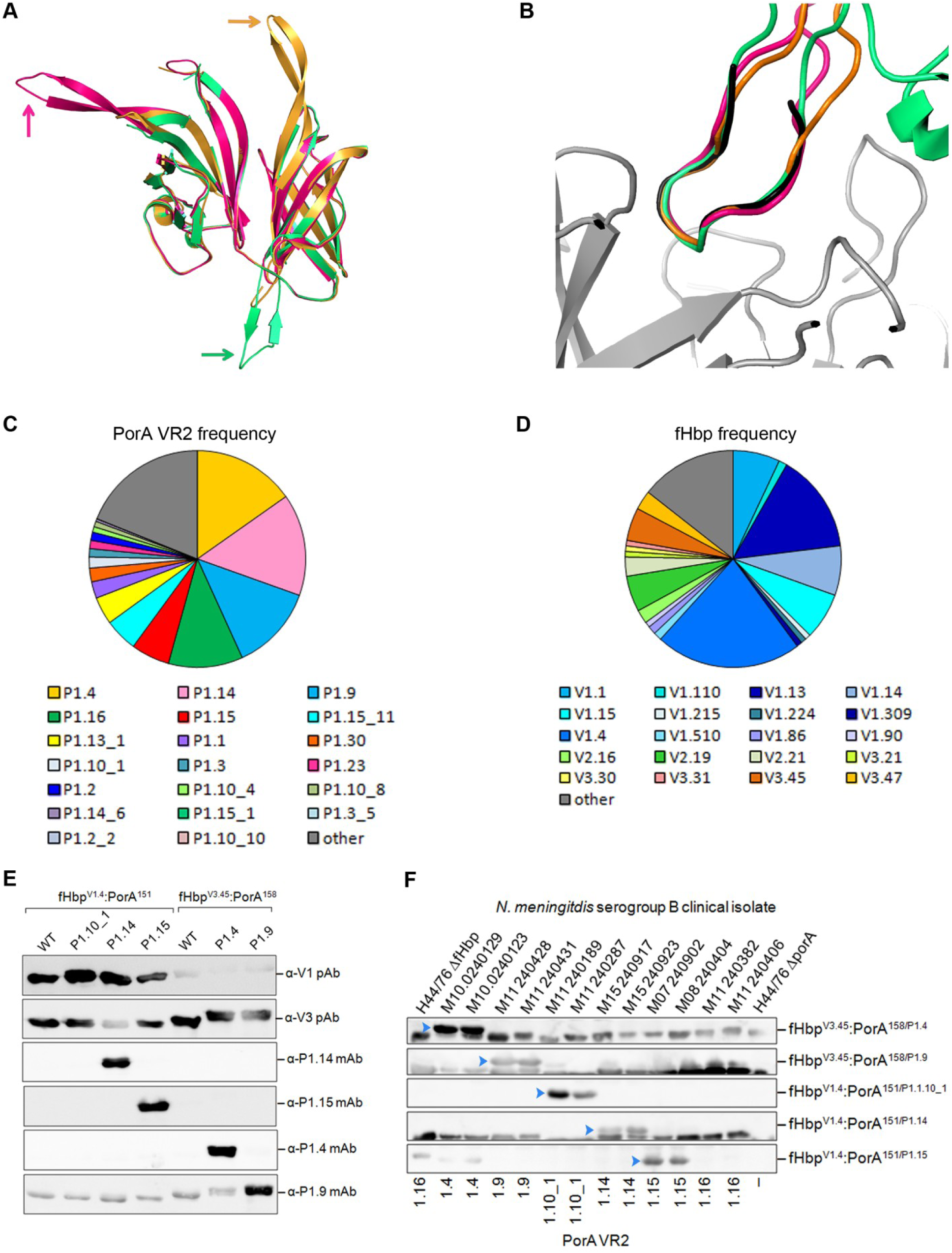
Structure of ChAs and frequency of fHbp/PorA alleles in the UK. (**A**) Alignment of fHbp scaffolds from fHbp^V1.4^:PorA^151/P1.16^ (pink), fHbp^V1.1^:PorA^294/P1.16^ (green) and fHbp^V1.4^:PorA^309/P1.16^ (orange), in each structure the P1.16 VR2 loop extends away from the main body of fHbp, this is indicated for each fHbp:PorA by the correspondingly coloured arrow. (**B**) Alignment of P1.16 VR2 region “KDTNNNL” from fHbp^V1.4^:PorA^151/P1.16^ (pink), fHbp^V1.1^:PorA^294/P1.16^ (green) and fHbp^V1.4^:PorA^309/P1.16^ (orange) with the P1.16 peptide “KDTNNNL” (black) in a complex with a bactericidal Fab fragment (grey) from mAb MN12H2 (PDB ID: 2MPA). Frequency of PorA VR2 (**C**) and fHbp variants (**D**) in *N. meningitidis* serogroup B strains (n=243) isolated in 2016 in the UK. Data downloaded from the Meningococcal Research Foundation, 27 June 2017. Other: remaining alleles that occur in <4 isolates. (**E**) Analysis of recombinant ChAs by SDS-PAGE and Western blot. Immunoblots are probed with α-PorA VR2 mAbs: P1.4, P1.9, P1.14 and P1.15. (**F**) Detection of PorA in a panel of *N. meningitidis* serogroup B isolates by mouse polyclonal antisera from ChAs fHbp^V1.4^:PorA^151/P1.1.10_1^, fHbp^V1.4^:PorA^151/P1.14^ and fHbp^V1.4^:PorA^151/P1.15^, fHbp^V3.45^:PorA^158/P1.4^ and fHbp^V3.45^:PorA^158/P1.9^.

### ChAs containing an expanded range of PorA VR2 loops generate immune responses

To test the adaptability of our fHbp:PorA ChAs, we generated several ChAs composed from different combinations of fHbp and PorA VR2. The comprehensive meningococcal genome data available for strains isolated in the UK enables construction of ChAs that have exact sequence matches to the most common antigens in a given region. In 2016, the most prevalent PorA VR2s in serogroup B *N. meningitidis* isolates were P1.4 (15.2 %), P1.14 (15.2 %), P1.9 (12.8 %), P1.16 (11.1 %) and P1.15 (5.8 %, **Figure 4C**). Whilst the most prevalent variant 1 and variant 3 fHbps were V1.4 and V3.45, present in 21.8 % and 4.9 % of serogroup B *N. meningitidis* isolates respectively (**Figure 4D**). We constructed five different ChAs, in which a PorA VR2 was inserted position 151 (V1.4) or position 158 (V3.45, **Figure 1B**). Following ChA expression and purification, Western blot analyses confirmed these ChAs all retained epitopes recognised by their cognate α-VR2 mAb and a-fHbp pAbs (**Figure 4E**).

To examine the ability of these fHbp:PorA ChAs to elicit immune responses, groups of CD1 mice were immunized with each ChA/alum (**Figure 2A**); antisera obtained post immunisation were pooled. To assess the resulting PorA immune responses, Western blot were conducted with pooled antisera and a panel of serogroup B *N. meningitidis* disease isolates. **Figure 4F** demonstrates that all ChAs elicited α-PorA antibodies that recognised their cognate PorA VR2. To evaluate α-PorA SBA responses, we performed SBAs with pooled ChA/alum antisera and serogroup B *N. meningitidis* strains with mismatched fHbp variants, to negate fHbp cross-protection. Titres range between ≥20 to ≥1280 and are above the ≥8 threshold for an accepted correlate of protective immunity against *N. meningitidis*(Andrews et al., 2003) (**Table 2**).

**Table 2:** Serum bactericidal assay titres

| Pooled antisera | Serogroup B isolate | fHbp variant | PorA VR2 | SBA titre |
| --- | --- | --- | --- | --- |
| fHbp<sup>V3.45</sup>:PorA<sup>158/P1.4</sup> | M10240123 | V1.92* | P1.4 | 1/160 |
| fHbp<sup>V3.45</sup>:PorA<sup>158/P1.9</sup> | M11240431 | V2.19 | P1.9 | 1/1280 |
| fHbp<sup>V1.4</sup>:PorA<sup>151/P1.1.10_1</sup> | M11240189 | V3.84 | P1.10_1 | 1/20 |
| fHbp<sup>V1.4</sup>:PorA<sup>151/P1.14</sup> | M15240853 | V3.45 | P1.14 | 1/640 |

α-PorA SBA titres generated using pooled ChA/alum antisera and serogroup B *N. meningitidis* isolates with mismatched fHbp variants. \* fHbp truncated at residue 242.

## Discussion

During infection pathogens present our immune system with an assortment of surface exposed lipid anchored and integral membrane proteins, both of which can be used as components in subunit vaccines. Whilst lipoproteins can be simply engineered for recombinant expression (by removal of their lipid anchor), integral membrane proteins present several challenges for vaccine development. Recombinant forms of integral membrane proteins are often poorly expressed and their native conformations may be compromised during purification, potentially reducing their ability to elicit immune responses against conformational epitopes, such as those found in surface loops (Bagal, Brown et al., 2013, Carpenter et al., 2008). Furthermore, immunisation with integral membrane proteins can generate irrelevant immune responses, which are directed towards epitopes masked by the outer membrane (Zhu, Thomas et al., 2005). To circumvent these issues, we used structure-based design to develop ChAs. We selected a key surface exposed epitope (VR2) from the integral membrane protein PorA and inserted it into the immunogenic scaffold of the lipoprotein fHbp. Multivalent ChAs generate immune responses against two key surface antigens that can elicit protective immunity (Claassen et al., 1996, Green et al., 2016, Kaaijk, van Straaten et al., 2013, Serruto et al., 2012), providing proof in principle that immunogenic epitopes from integral membrane proteins can be introduced into soluble molecular scaffolds to create ChAs.

Combining two antigens within a single recombinant ChA could diminish the immunogenicity of one or both antigens. We found that fHbp in all ChAs is highly immunogenic, inducing significant bactericidal antibody responses, with titres in excess of those correlated with protection (Andrews et al., 2003). In addition, the PorA VR2 loop in ChAs can induce antibody responses even when presented away from its native environment in the outer membrane. While all ChAs induced antibodies that detect PorA on the surface of *N. meningitidis*, not all induced bactericidal PorA antibodies. We observed a mixed bactericidal PorA response, which depended on the PorA VR2 variant, the position of PorA VR2 in the fHbp scaffold and the adjuvant used for immunisation, indicating that several parameters are crucial for obtaining bactericidal responses.

Linear PorA VR2 P1.16 peptides elicit antibodies that fail to recognise the native protein and are non-bactericidal, while cyclic PorA VR2 peptides, with identical residues but fixed into a β-turn, can elicit antibodies that recognise native PorA and are bactericidal (Christodoulides, McGuinness et al., 1993, Christodoulides, Rattue et al., 1999, Hoogerhout, Donders et al., 1995, Oomen et al., 2003, Oomen et al., 2005). Structural data shows that when the structure of cyclic VR2 peptide mirrors that of a linear peptide bound by a bactericidal mAb, and thus locked into an immunogenic conformation, the cyclic peptide induces bactericidal responses (Oomen et al., 2003, Oomen et al., 2005). In ChAs, the N-and C-termini of the VR2 loop are bound by neighbouring fHbp β-strands, fixing the VR2 epitope into a β-turn, as observed in cyclic peptides. This was confirmed by the atomic structures of ChAs, in which the PorA adopts the same conformation as when bound by a bactericidal Fab fragment (van den Elsen et al., 1997). In previous work, cyclic peptides were coupled to carrier proteins and used with adjuvants not licensed for human use (Christodoulides et al., 1993, Christodoulides et al., 1999, Oomen et al., 2003, Oomen et al., 2005). In the resulting SBAs, titres were only observed with antisera from some mice (Oomen et al., 2005), similar to our findings with ChAs and licensed adjuvants.

Alum allows extended antigen presentation and stimulates T-helper (Th)−2 responses, predominantly producing IgG1 and IgE (Marrack, McKee et al., 2009, Moingeon, Haensler et al., 2001, Petrovsky & Aguilar, 2004), while MPLA typically enhances Th1 responses, inducing IgG2a, IgG2b, and IgG3 (Baker, Hiernaux et al., 1988, Germann, Bongartz et al., 1995). Murine IgG2a has a greater ability than murine IgG1 to activate the classical pathway (Leatherbarrow & Dwek, 1984, Michaelsen, Kolberg et al., 2004). Consistent with this, immunisation with ChA/MPLA resulted in significant α-fHbp IgG2a responses, leading to higher α-fHbp SBA titres than observed with ChA/Alum. Of note, antisera raised against fHbp^V1.1^:PorA^309/P1.16^ /MPLA had α-PorA SBA titre and contained significant levels of α-PorA IgG2a and IgM.

Using structure-based design, we generated ChAs that retain epitopes of fHbp and PorA and generate immune responses against both antigens. Our work demonstrates that a soluble antigen can be exploited as a scaffold to display epitopes from an integral membrane protein. The development of ChAs paves the way for exploiting immunogenic, but difficult to express, membrane proteins in vaccines; a valuable approach for vaccine design that could be applied to other pathogens. Furthermore, we generated ChAs with exact sequence matches to the most prevalent fHbp and PorA antigens expressed by serogroup B *N. meningitidis* strains currently circulating in the UK; thereby demonstrating pathogen antigenic diversity can be circumvented by tailoring ChA composition to match the prevalent antigens. A vaccine composed of the three most common fHbps from each variant group (fHbps V1.4, V2.19 and V3.45) with a single PorA VR2 insertion (PorA VR2 P1.4, P1.9 and P1.14) would give exact sequence coverage against 57% of serogroup B strains circulating in the UK in 2016, which compares favourably with the currently licensed meningococcal serogroup B vaccines (Green et al., 2016, Serruto et al., 2012).

## Materials and Methods

### Bacterial strains and growth

The bacterial strains used in this study are shown in **Table S2** and **Table S3**. *N. meningitidis* was grown in the presence of 5% CO_2_ at 37°C on Brain Heart Infusion (BHI, Oxoid, Basingstoke, United Kingdom) plates with 5% (v/v) horse serum (Oxoid) at 37°C. *Escherichia coli* was grown on Luria-Bertani (LB) agar plates or LB liquid at 37°C supplemented with 100 μg μl^−1^ carbenicillin.

### Expression and purification of fHbp-PorAs

N-terminally truncated V1.1 fHbp was amplified from MC58 genomic DNA using primers fHbp F1 and fHbp R4 (**Table S4**). The PorA P1.16 VR2 (YYTKDTNNNLTLV) was introduced into one of six positions in fHbp by overlap PCR using the primers in **Table S4**. The PorA VR2 loops P1.4 (HVVVNNKVATHVP), P1.9 (YVDEQSKYHA), P1.10_1 (HFVQNKQNQPPTLVP), P1.14 (YVDEKKMVHA) and P1.15 (HYTRQNNADVFVP) were introduced into a single position in fHbp by overlap PCR using the primers in **Table S4**. PCR products were digested with *NdeI* and *XhoI* (NEB) then ligated into pET21b (Novagen); constructs were confirmed by sequencing. Protein expression was performed in *E. coli* strain B834. Expression cultures were incubated at 37°C, upon reaching an OD_600_ of ~0.8, protein expression was induced with 1 mM IPTG. Cultures were harvested after overnight expression at 37°C. Bacteria were re-suspended in Buffer A (50 mM Na-phosphate pH 8.0, 300 mM NaCl, 30 mM Imidazole) and purified by Nickel affinity chromatography (His-trap FF Crude, GE Healthcare) at room temperature. Columns were washed with 25 column volumes (CV) of Buffer A, then 20 CV 80:20 Buffer A: Buffer B (50 mM Na-phosphate pH 8.0, 300 mM NaCl, 300 mM Imidazole), elution was performed with 10 CV 40:60 Buffer A: Buffer B. Proteins were dialysed overnight at 4°C into 50 mM Na-Acetate pH 5.5 buffer and further purification was achieved by ion exchange chromatography (HiTrapSP HP, GE Healthcare) at room temperature with a 0-1 M NaCl gradient in 50 mM Na-Acetate pH 5.5 buffer, followed by gel filtration using a HiLoad 16/600 Superdex 75 pg (GE Healthcare) column equilibrated with phosphate buffered saline (PBS, Oxoid).

### Generation of N. meningitidis strains

Deletion of *porA* was performed by replacing the open reading frame with a tetracyline resistance cassette. Briefly, ~500 bp of the up-and downstream regions of the *porA* locus flanking a tetracycline resistance cassette was generated by PCR using primers PorA KO: F1, R1, F2, R2 (**Table S4**) and the mega-primer method (Ke & Madison, 1997). The PCR product was transformed into *N. meningitidis* H44/76 and H44/76Δ*fHbp* as described previously (Exley, Shaw et al., 2005). Genomic DNA was obtained (Wizard^®^ genomic DNA purification kit) from the resulting strains and both mutations were backcrossed into the WT H44/76 background using genomic DNA (Exley et al., 2005).

### Western blot analyses

Western blots of purified proteins (0.5 μg) or 10 μl of *N. meningitidis* cell lysate (Jongerius, Lavender et al., 2013) were probed with one of the following primary sera: PorA P1.4 mAb (NIBSC cat: 02/148, diluted 1 in 500), PorA P1.9 mAb (NIBSC cat: 05/190, diluted 1 in 250), PorA P1.14 mAb (NIBSC cat: 03/142, diluted 1 in 500), PorA P1.15 mAb (NIBSC cat: 02/144, diluted 1 in 1,000), PorA P1.16 mAb (NIBSC cat: 01/538, diluted 1 in 1,000), pAb to fHbp V1.1 (Jongerius et al., 2013) (diluted 1 in 1,000), or sera from mice immunised with an fHbp:PorA ChA (diluted 1 in 500). Following incubation with anti-mouse HRP conjugated secondary antibodies (diluted 1 in 10,000), membranes were visualised via ECL (GE Healthcare) on an LAS-4000 (FujiFilm).

### Generation of immune sera

Immunisations were performed with each fHbp:PorA ChA and PBS controls, using alum or MPLA as the adjuvant. Alum immunisations were prepared by incubating 20 μg fHbp:PorA or PBS, 2 % Alhydrogel (Invivogen), 10 mM Histidine-HCl pH 6.5 and 155 mM NaCl overnight at 4oC on an end-over-end rocker. For MPLA immunisations, lyophilised MPLA (Invivogen) was resuspended in sterile H_2_O by incubating for 5 minutes in a sonicating water bath. 10 μg of MPLA was mixed at room temperature with 20 μg fHbp:PorA or PBS, 10 mM Histidine-HCl pH 6.5 and 155 mM NaCl. Groups of eight female CD1 mice (~6 weeks old, Charles Rivers, Margate) were immunised with three intraperitioneal injections administered on days 0, 21 and 35. Sera was obtained on day 49 following cardiac puncture under terminal anaesthesia. All procedures were conducted in accordance with UK Home Office guidelines. All sera were stored at −80°C until required and once defrosted sera were stored at 4°C.

### Serum Bactericidal Assays

SBAs were performed as previously described (Jongerius et al., 2013), with the following modifications. *N. meningitidis* was suspended in Dubecco’s PBS with cations (Gibco) supplemented with 0.1 % glucose (DPBS-G) to a final concentration of 1.25 x 10^4^ CFU ml^−1^. Baby rabbit complement (Cedar lane, lot #15027680) was diluted with DPBS-G to a final dilution of 1 in 10. Serum, pooled or from individual mice, was heat inactivated for one hour at 56°C and added to the wells in a serial two-fold dilution, starting with a dilution of 1 in 5 or higher. Control wells contained no serum or no complement. Following static incubation for one hour at 37°C in the presence of 5% CO_2_, 10 µl from each well was plated onto BHI plates in triplicate and colonies from surviving bacteria counted. The bactericidal activity is expressed as the dilution of serum required to kill ≥ 50% of bacteria in assays containing both complement and serum in comparison with control assays containing serum or complement alone. SBAs using pooled sera were repeated three times, and assays using sera from individual mice were repeated twice. SBA titres were input into GraphPad Prism and statistical analyses comparing titres obtained from alum immunisations with titres obtained from MPLA immunisations were performed using two-way ANOVA (statistical significance of *p* ≤ 0.05) and Dunnett’s method of multiple comparisons.

### Surface Plasmon Resonance

SPR was performed using a Biacore 3000 (GE Healthcare). Recombinant ChAs were dissolved in 50 mM sodium acetate pH 4.5 and immobilized on a CM5 sensor chip (GE Healthcare). Increasing concentrations of human complement FH domains 6 and 7 (1 nM-16 nM) were injected over the flow channels at 40 µl min^−1^). Dissociation was allowed for 300 seconds. BIAevaluation software was used to calculate the *K*_D_

### Structural biology

fHbp:PorAs in PBS were concentrated to 9 mg ml^−1^ and screened for crystal formation via the sitting drop method at a 0.4:0.6 ratio of protein to mother liquor. Crystals of chimeras fHbp^V1.4^:PorA^151/P1.16^, fHbp^V1.1^:PorA^294/P1.16^, fHbp^V1.1^:PorA^309/P1.16^ and fHbp^V1.4^:PorA^309/P1.16^ grew under the respective conditions: 2.0 M ammonium sulphate, 0.15 M sodium citrate pH 5.5; 20% (w/v) PEG4000, 0.3 M ammonium sulphate; 0.01 M zinc chloride, 0.1 M sodium acetate, 20% (w/v) PEG6000; and 0.2 M potassium formate, 20% (w/v) PEG3350 respectively. Crystals were transferred to a cryoprotectant solution comprised of mother liquor and 30% ethylene glycol. Data were collected at Diamond light sourceand integrated/scaled using the Xia2 programme. Molecular replacement was carried out using Phaser from the CCP4 package (Winn, Ballard et al., 2011), with the V1.1 fHbp structure used as a search model (PDB 2W80). Refmac5 (Murshudov, Vagin et al., 1997), Coot (Emsley & Cowtan, 2004) and Phenix (Afonine, Grosse-Kunstleve et al., 2013, Afonine, Grosse-Kunstleve et al., 2012, Afonine, Grosse-Kunstleve et al., 2009, Headd, Echols et al., 2012) were utilised for rebuilding and refinement, with structural validation performed in Molprobity (Chen, Arendall et al., 2010). Refinement statistics for each structure are detailed in **Table S5**. PDB accession codes are: 5NQP, fHbp^V1.4^:PorA^151/P1.16^; 5NQX, fHbp^V1.1^:PorA^294/P1.16^_;_ 5NQZ, fHbp^V1.1^:PorA^309/P1.16^; 5NQY, fHbp^V1.4^:PorA^309/P1.16^.

### Differential scanning calorimetry

20 μM of purified protein in PBS was subjected to a 20 to 120°C temperature gradient on a Malvern VP Capillary DSC. Melting temperature (T_m_) is recorded for the fHbp N-terminal and C-terminal β-barrels.

### Flow cytometry

*N. meningitidis* strains H44/76, H44/76Δ*fHbp*, H44/76Δ*porA* and H44/76Δ*fHbp*Δ*porA* (1x10^9^ cells) were fixed in 3% paraformaldehyde (PFA) for one hour. PFA was removed by washing cells three times with PBS. To measure binding of serum antibodies to fHbp and PorA, 1x10^8^ cells were incubated for one hour at 4°C, shaking at 1,250 rpm, with sera diluted 1 in 50 in PBS. Cells were washed three times with PBS, then incubated for 30 minutes at 4°C with shaking at 1,250 rpm, with a secondary antibody (all ThermoFisher Scientific). Final dilutions of secondary antibodies are as follows: 5 μg ml^−1^ for Alexa-488 conjugated anti-mouse IgAGM, and 2.5 μg ml^−1^ for each of Alexa-488 conjugated anti-mouse IgM, IgG3 and IgG2b, and Alexa-647 conjugated anti-mouse IgG2a and IgG1. Cells were washed three times with PBS and then resuspended in PBS for analysis on the FACSCalibur (BD Biosciences). Data were imported into the FlowJo^TM^ data analysis package and transformed to the Logicle scale. A uniform gate was applied to all data sets, based on the *N. meningitidis* population visualised by forward and side-scatter, and the geometric mean of fluorescence calculated for FL1H (Alexa-488 signal) and FL4H (Alexa-647 signal). Each flow experiment was repeated three times, and the FL1H and FL4H geometric means were input into Graphpad Prism. Two-way ANOVA (statistical significance of *p* ≤ 0.05, using Dunnett’s method of multiple comparison) was used to compare the Alexa-488 or Alexa-647 geometric means with control Alexa-488 or Alexa-647 geometric means. Control results were obtained by incubating PFA fixed cells with sera from mice immunised with PBS and Alum or MPLA adjuvants, and the corresponding Alexa-fluor labelled secondary Ig. Two-way ANOVA multiple-comparisons were performed with geometric means from sets of experiments using the same secondary antibody, *N. meningitidis* strain and adjuvant; the only variable factor is the fHbp:PorA used for immunisation.

## Acknowledgments

We gratefully acknowledge our financial support from Action Medical Research (Award number GN2205), Wellcome Trust Senior Investigator Awards (Award numbers

102908/Z/13/Z and 100298/Z/13/Z) and Medical Research Council (Award number MR/M011984/1). We thank David Staunton for performing the DSC analysis, Sophie Andrews and Sasha Burgess for assisting with the production of expression vectors, Diamond and ESRF synchotrons for provision of beam time, and the Diamond-I02, Diamond-I04 and ESRF-ID29 beamline staff for help with beamline preparation and data collection. We gratefully thank the Meningococcal Reference Unit, Public Health England, for the meningococcal serogroup B disease isolates.

## Author contributions

SH and IJ designed and performed experiments. CMT, SJ, SML and RME designed experiments. SJ and SML performed structural studies. CMT managed the project. SH prepared the figures. SH, CMT, IJ, RME, SJ and SML wrote the manuscript.

## Supplementary information

**Figure S1.**
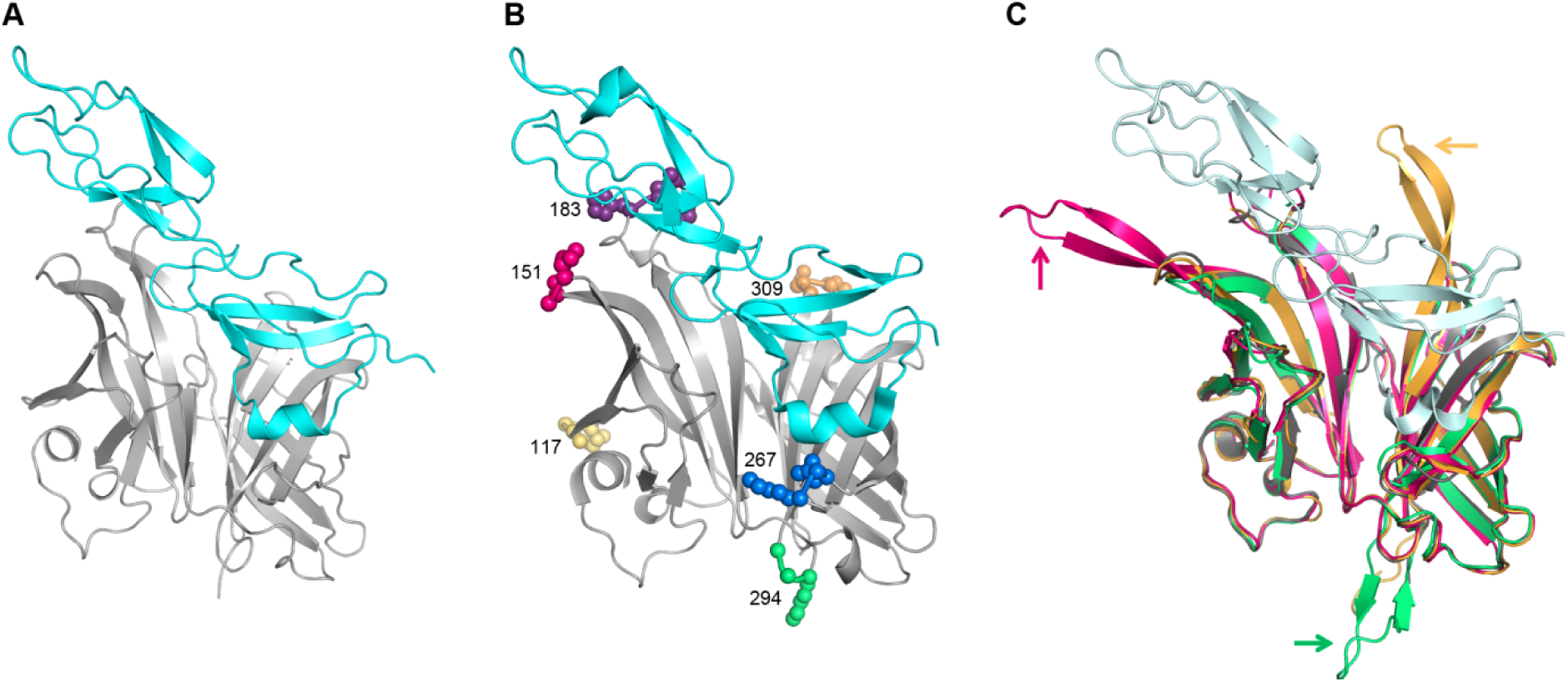
Structural alignments of ChAs with wild type fHbp and PorA VR2 peptide. (**A**) Complex formed by V1.1 fHbp (grey) and human complement FH domains 6 and 7 (cyan, PDB ID 2W80). (**B**) Locations of the six fHbp residues replaced with the PorA P1.16 VR2 loop in relation to the FH binding interface. (**C**) Secondary structure alignment of fHbp:PorAs with V1.1 fHbp (grey) in a complex with human complement FH domains 6 and 7 (light blue); fHbp^v1.4^:PorA^151/P1.16^ (pink), fHbp^v1.1^:PorA^294/P1.16^ (green), fHbp^v1.4^:PorA^309/P1.16^ (orange). PorA P1.16 VR2 loops are indicated by the correspondingly coloured arrows.

**Figure S2.**
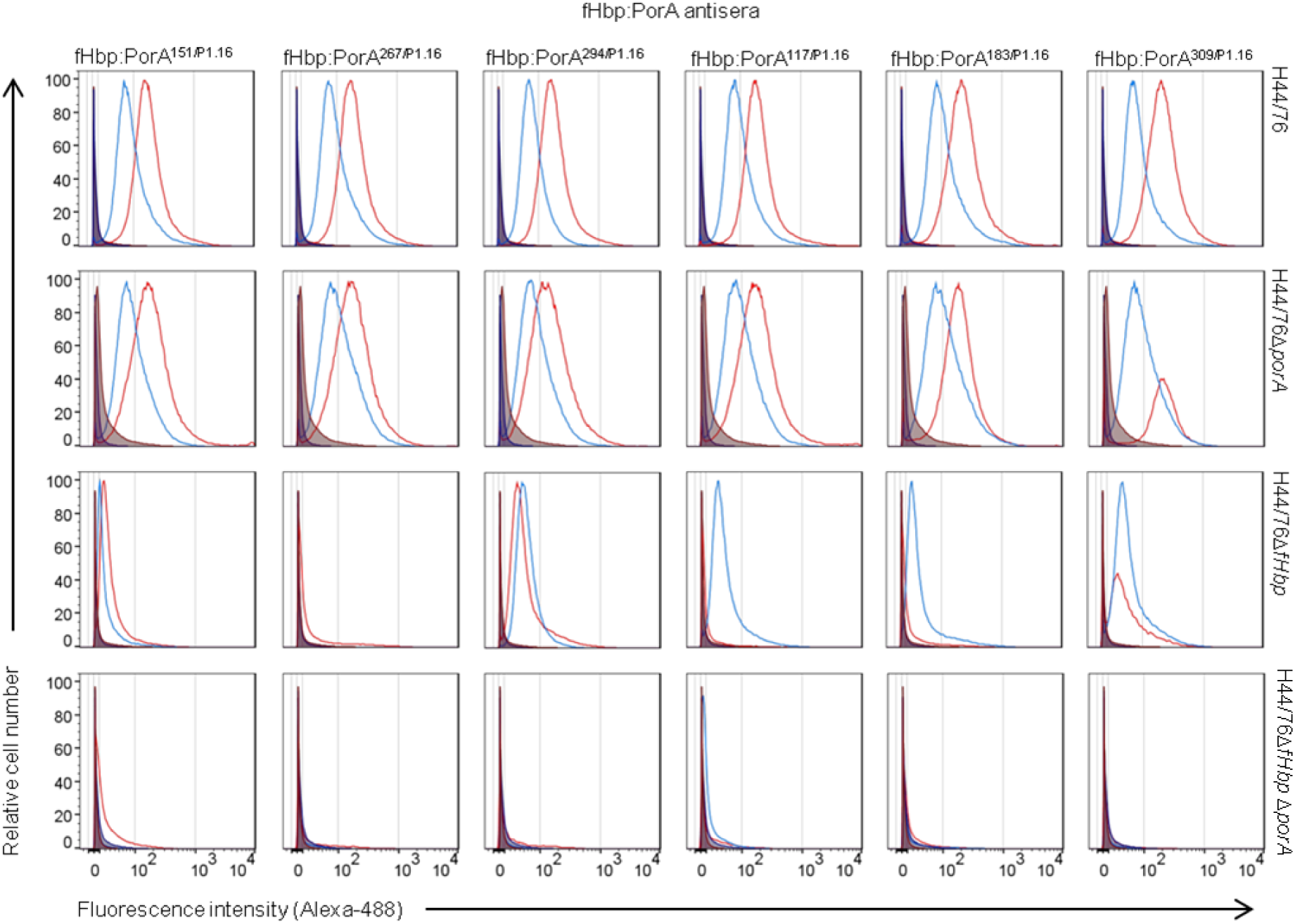
Flow cytometry histograms showing detection of fHbp and PorA by ChA antisera. Histograms of flow cytometry experiments conducted with alum (blue) and MPLA (red) ChA antisera binding to fixed *N. meningitidis* H44/76 strains: wild type (WT), Δ*fHbp*, Δ*porA* and Δ*fHbp*Δ*porA. fHbp* and PorA antibodies bound to *N. meningitidis* were detected using Alexa-488 conjugated IgA/G/M secondary antibody. Alum PBS and MPLA PBS antisera are shown in dark blue and dark red shaded profiles, and secondary IgAGM control is shown in black. Comparison of the geometric means from these histograms, and the histograms from two other independent repeats, is shown in Figures 2C and 2D

**Table S1:** Nomenclature of fHbp:PorA ChAs

| fHbp scaffold | PorA VR2 loop inserted | Residue position of PorA VR2 loop insertion | ChA |
| --- | --- | --- | --- |
| V1.1 | P1.16 | 151 | fHbp<sup>V1.1</sup>:PorA<sup>151/P1.16</sup> |
| V1.1 | P1.16 | 267 | fHbp<sup>V1.1</sup>:PorA<sup>267/P1.16</sup> |
| V1.1 | P1.16 | 294 | fHbp<sup>V1.1</sup>:PorA<sup>294/P1.16</sup> |
| V1.1 | P1.16 | 117 | fHbp<sup>V1.1</sup>:PorA<sup>117/P1.16</sup> |
| V1.1 | P1.16 | 183 | fHbp<sup>V1.1</sup>:PorA<sup>183/P1.16</sup> |
| V1.1 | P1.16 | 309 | fHbp<sup>V1.1</sup>:PorA<sup>309/P1.16</sup> |
| V1.4 | P1.16 | 151 | fHbp<sup>V1.4</sup>:PorA<sup>151/P1.16</sup> |
| V1.4 | P1.16 | 309 | fHbp<sup>V1.4</sup>:PorA<sup>309/P1.16</sup> |
| V1.4 | P1.10_1 | 151 | fHbp<sup>V1.4</sup>:PorA<sup>151/P1.1.10_1</sup> |
| V1.4 | P1.14 | 151 | fHbp<sup>V1.4</sup>:PorA<sup>151/P1.14</sup> |
| V1.4 | P1.15 | 151 | fHbp<sup>V1.4</sup>:PorA<sup>151/P1.15</sup> |
| V3.45 | P1.4 | 158 | fHbp<sup>V3.45</sup>:PorA<sup>158/P1.4</sup> |
| V3.45 | P1.9 | 158 | fHbp<sup>V3.45</sup>:PorA<sup>158/P1.9</sup> |

**Table S2:**
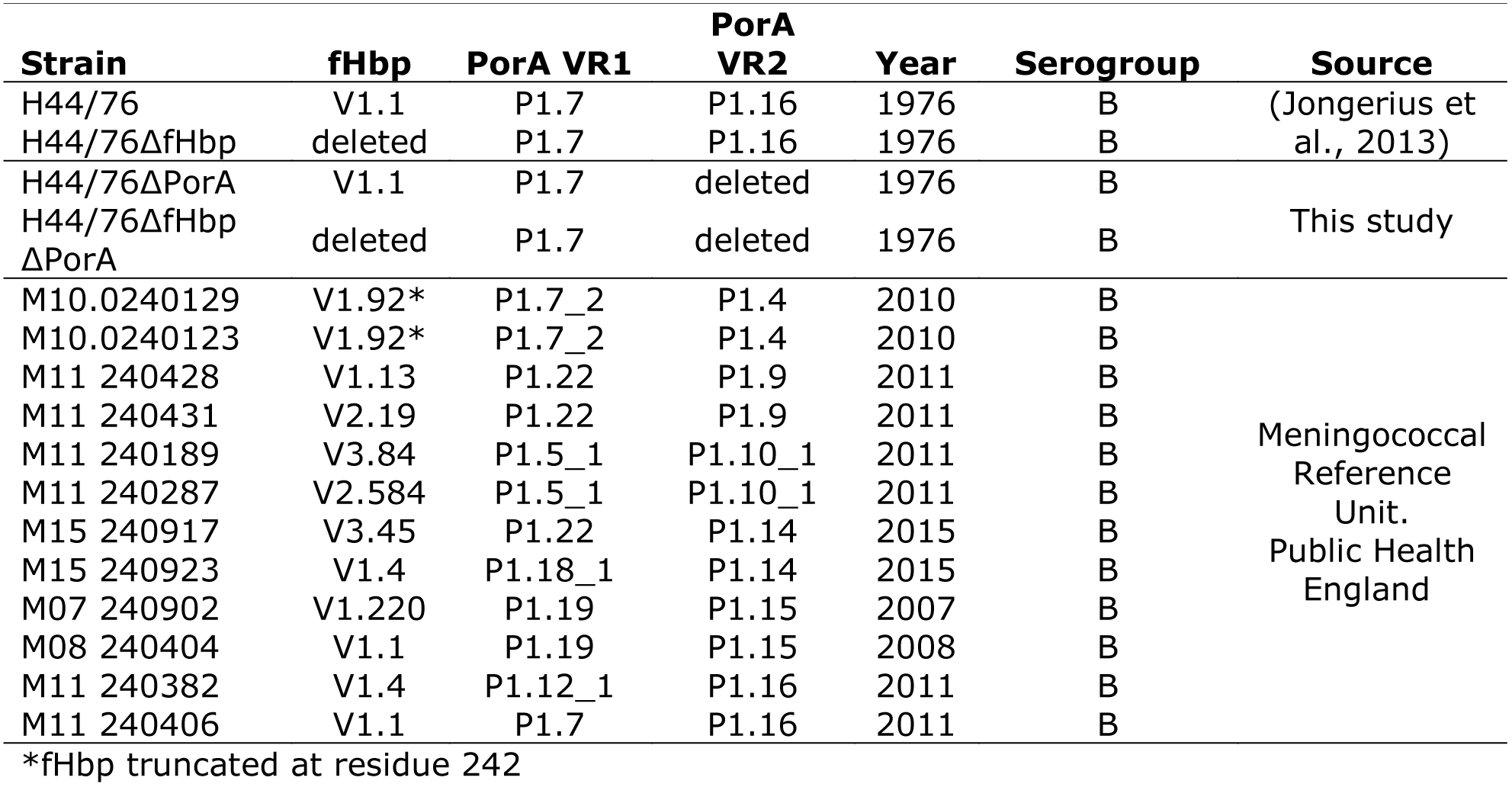
Neisseria meningitidis strains

**Table S3:** Escherchia coli strains

| **Strain** | **Plasmid** | **Protein expressed** |
| --- | --- | --- |
| B834 | pET21b-fHbp<sup>V1.1</sup> | V1.1 fHbp |
| B834 | pET21b-fHbp<sup>V1.1</sup>:PorA<sup>151/P1.16</sup> | fHbp<sup>V1.1</sup>:PorA<sup>151/P1.16</sup> |
| B834 | pET21b-fHbp<sup>V1.1</sup>:PorA<sup>267/P1.16</sup> | fHbp<sup>V1.1</sup>:PorA<sup>267/P1.16</sup> |
| B834 | pET21b-fHbp<sup>V1.1</sup>:PorA<sup>294/P1.16</sup> | fHbp<sup>V1.1</sup>:PorA<sup>294/P1.16</sup> |
| B834 | pET21b-fHbp<sup>V1.1</sup>:PorA<sup>117/P1.16</sup> | fHbp<sup>V1.1</sup>:PorA<sup>117/P1.16</sup> |
| B834 | pET21b-fHbp<sup>V1.1</sup>:PorA<sup>183/P1.16</sup> | fHbp<sup>V1.1</sup>:PorA<sup>183/P1.16</sup> |
| B834 | pET21b-fHbp<sup>V1.1</sup>:PorA<sup>309/P1.16</sup> | fHbp<sup>V1.1</sup>:PorA<sup>309/P1.16</sup> |
| B834 | pET21b-fHbp<sup>V1.4</sup>:PorA<sup>151/P1.16</sup> | fHbp<sup>V1.4</sup>:PorA<sup>151/P1.16</sup> |
| B834 | pET21b-fHbp<sup>V1.4</sup>:PorA<sup>309/P1.16</sup> | fHbp<sup>V1.4</sup>:PorA<sup>309/P1.16</sup> |
| B834 | pET21b-fHbp<sup>V1.4</sup>:PorA<sup>151/P1.1.10_1</sup> | fHbp<sup>V1.4</sup>:PorA<sup>151/P1.1.10_1</sup> |
| B834 | pET21b-fHbp<sup>V1.4</sup>:PorA<sup>151/P1.14</sup> | fHbp<sup>V1.4</sup>:PorA<sup>151/P1.14</sup> |
| B834 | pET21b-fHbp<sup>V1.4</sup>:PorA<sup>151/P1.15</sup> | fHbp<sup>V1.4</sup>:PorA<sup>151/P1.15</sup> |
| B834 | pET21b-fHbp<sup>V3.45</sup>:PorA<sup>158/P1.4</sup> | fHbp<sup>V3.45</sup>:PorA<sup>158/P1.4</sup> |
| B834 | pET21b-fHbp<sup>V3.45</sup>:PorA<sup>158/P1.9</sup> | fHbp<sup>V3.45</sup>:PorA<sup>158/P1.9</sup> |

**Table S4:**
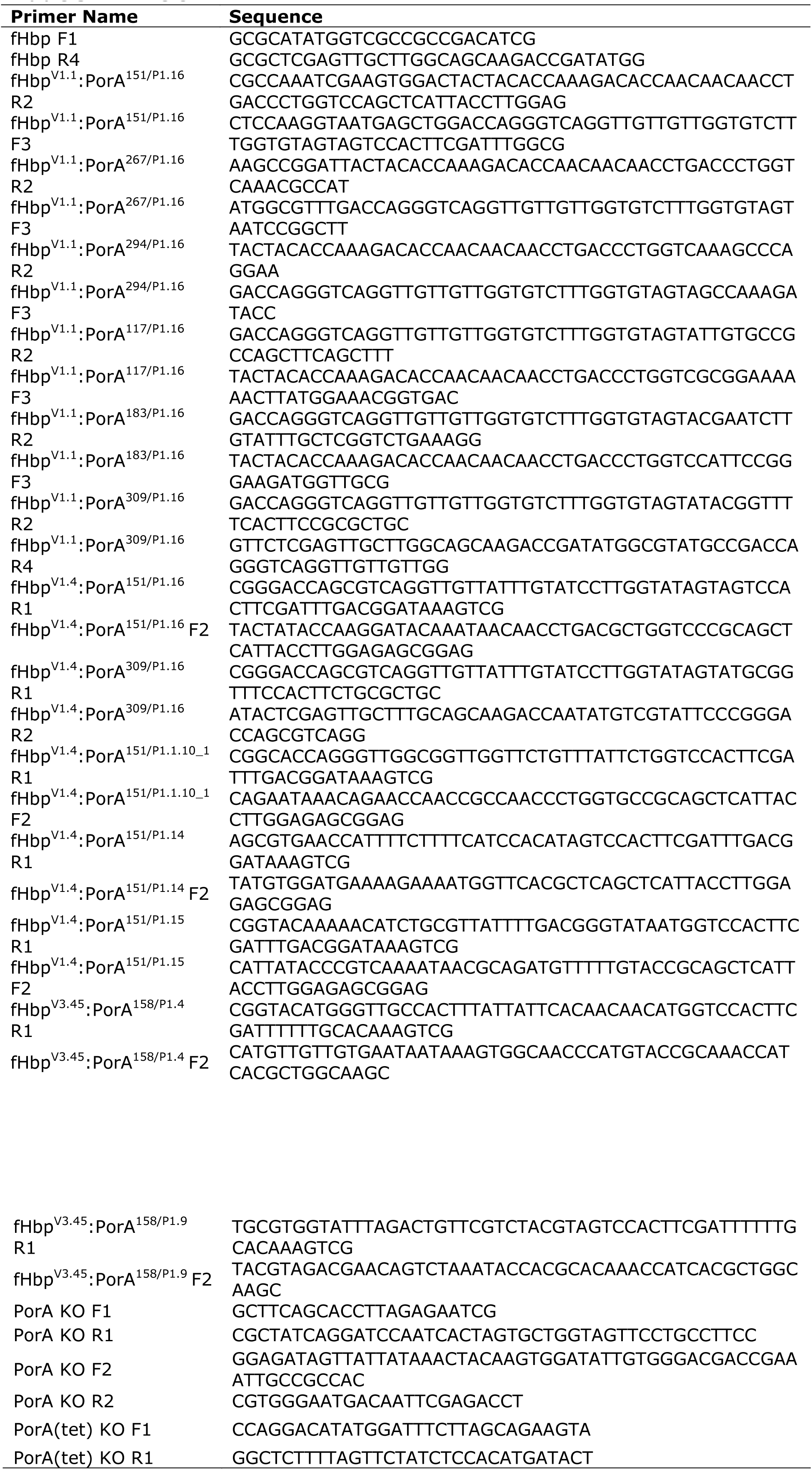
Primers

**Table S5:** Data collection and refinement statistics.

| Protein | fHbp<sup>V1.4</sup>:PorA<sup>151/P</sup><br>1.16 | fHbp<sup>V1.1</sup>:PorA<sup>294</sup><br>/P1.16 | fHbp<sup>V1.4</sup>:PorA<sup>309/P</sup><br>1.16 | fHbp<sup>V1.1</sup>:PorA<sup>309</sup><br>/P1.16 |
| --- | --- | --- | --- | --- |
| PDB Code | 5nqp | 5nqx | 5nqy | 5nqz |
| **Data collection statistics** |  |  |  |  |
| Beamline | Diamond I02 | Diamond I02 | ESRF ID29 | Diamond I04-1 |
| Wavelength (Å) | 0.97949 | 0.97858 | 0.97623 | 0.92818 |
| Resolution limits (Å)<sup>a</sup> | 40.74-2.86<br>(2.93-2.86) | 81.65-3.66<br>(3.76-3.66) | 51.17-2.60<br>(2.67-2.60) | 58.55-1.63<br>(1.67-1.63) |
| Space group | C222<sub>1</sub> | P22<sub>1</sub>2<sub>1</sub> | P2<sub>1</sub>2<sub>1</sub>2<sub>1</sub> | C2 |
| Unit cell dimensions (Å, °) | 100.7,171.4,131.5<br>90,90,90 | 94.6,114.9,160.3<br>90,90,90 | 56.9,62.6,88.7<br>90,90,90 | 91.4,76.3,83.9<br>90,91.3,90 |
| Unique reflections | 26403<br>(1923) | 19820<br>(1431) | 10208<br>(743) | 70772<br>(4967) |
| Multiplicity<sup>a</sup> | 4.5<br>(4.4) | 4.4<br>(4.6) | 6.4<br>(6.6) | 3.0<br>(2.2) |
| Completeness (%)<sup>a</sup> | 99.1<br>(99.4) | 99.3<br>(99.9) | 99.6<br>(99.4) | 98.6<br>(93.9) |
| *I*/$\sigma$(*I*)<sup>a</sup> | 17.1<br>(2.0) | 7.6<br>(1.9) | 16.7<br>(2.3) | 12.7<br>(1.5) |
| *R*<sub>merge</sub> (%)<sup>a,b</sup> | 0.074<br>(0.645) | 0.187<br>(0.833) | 0.073<br>(0.845) | 0.062<br>(0.615) |
| *R*<sub>pim</sub> (%) | 0.038<br>(0.340) | 0.101<br>(0.439) | 0.032<br>(0.356) | 0.041<br>(0.534) |
| **Refinement statistics** |  |  |  |  |
| Resolution limits (Å) | 40.74-2.86<br>(2.97-2.86) | 81.65-3.66<br>(3.85-3.66) | 51.17-2.60<br>(2.86-2.60) | 58.55-1.63<br>(1.67-1.63) |
| Number of reflections in working set | 25137<br>(2746) | 18807<br>(2650) | 9665<br>(2369) | 63880<br>(4735) |
| Number of reflections in test set | 1238<br>(134) | 975<br>(121) | 498<br>(120) | 3445<br>(225) |
| $R$ factor of working set<sup>a,c</sup> | 0.198<br>(0.303) | 0.232<br>(0.321) | 0.226<br>(0.351) | 0.162<br>(0.289) |
| $R_{\text{free}}$<sup>a,d</sup> | 0.239<br>(0.351) | 0.282<br>(0.368) | 0.271<br>(0.420) | 0.194<br>(0.297) |
| Number of atoms (protein/water/other) | 5895/28/84 | 9572/0/0 | 1994/4/4 | 4046/540/45 |
| Residues in Ramachandran favoured region (%) | 95.2 | 97.8 | 95.3 | 97.0 |
| Ramachandran outliers (%) | 0.1 | 0.1 | 0.0 | 0.2 |
| r.m.s.d. bond lengths (Å) | 0.002 | 0.002 | 0.002 | 0.02 |
| r.m.s.d. bond angles (°) | 0.584 | 0.424 | 0.449 | 1.98 |

<sup>a</sup>Numbers in parentheses refer to the appropriate outer shell.
<sup>b</sup> $R_{\text{merge}} = 100 \times (\sum_{hkl} \sum_i |I(hkl;i) - \langle I(hkl) \rangle| / \sum_{hkl} \sum_i I(hkl;i))$ , where $I(hkl;i)$ is the intensity of an individual measurement of a reflection and $\langle I(hkl) \rangle$ is the average intensity of that reflection.
<sup>c</sup> $R_{\text{factor}} = (\sum_{hkl} ||F_{\text{obs}}| - |F_{\text{calc}}|| / \sum_{hkl} |F_{\text{obs}}|)$ , where $|F_{\text{obs}}|$ and $|F_{\text{calc}}|$ are the observed and calculated structure factor amplitudes.
<sup>d</sup> $R_{\text{free}}$ equals the $R$ -factor of test set (5% of the data removed prior to refinement).
r.m.s.d.: root mean square deviation from ideal geometry.

## References

GraphPad Prism version 6.00 for Windows. In www.graphpad.com: GraphPad Software, La Jolla California USA

Meningitis Research Foundation Meningococcus Genome Library developed by Public health England, the Wellcome Trust Sanger Institute and the University of Oxford as a collaboration. In http://www.meningitis.org/research/genome: Meningitis Research Foundation

(2013) Gonococcal resistance to antimicrobials surveillance programme (GRASP) action plan for England and Wales: informing the public health response. In http://webarchive.nationalarchives.gov.uk/20140714084352/http://www.hpa.org.uk/webc/HPAwebFile/HPAweb_C/1317138215954: Health Protection Agency

(2016) Surveillance of antimicrobial resistance in Neisseria gonorrhoeae Key findings from the Gonococcal Resistance to Antimicrobials Surveillance Programme (GRASP). In https://www.gov.uk/government/uploads/system/uploads/attachment_data/file/567602/GRASP_Report_2016.pdf: Public Health England

Afonine PV, Grosse-Kunstleve RW, Adams PD, Urzhumtsev A (2013) Bulk-solvent and overall scaling revisited: faster calculations, improved results. Acta Crystallogr D Biol Crystallogr 69: 625–34

Afonine PV, Grosse-Kunstleve RW, Echols N, Headd JJ, Moriarty NW, Mustyakimov M, Terwilliger TC, Urzhumtsev A, Zwart PH, Adams PD (2012) Towards automated crystallographic structure refinement with phenix.refine. Acta Crystallogr D Biol Crystallogr 68: 352–67

Afonine PV, Grosse-Kunstleve RW, Urzhumtsev A, Adams PD (2009) Automatic multiple-zone rigid-body refinement with a large convergence radius. J Appl Crystallogr 42: 607–615

Andre FE, Booy R, Bock HL, Clemens J, Datta SK, John TJ, Lee BW, Lolekha S, Peltola H, Ruff TA, Santosham M, Schmitt HJ (2008) aVaccination greatly reduces disease, disability, death and inequity worldwide. Bull World Health Organ 86: 140–6

Andrews N, Borrow R, Miller E (2003) Validation of serological correlate of protection for meningococcal C conjugate vaccine by using efficacy estimates from postlicensure surveillance in England. Clin Diagn Lab Immunol 10: 780–6

Arístegui J, Usonis V, Coovadia H, Riedemann S, Win KM, Gatchalian S, Bock HL (2003) Facilitating the WHO expanded program of immunization: the clinical profile of a combined diphtheria, tetanus, pertussis, hepatitis B and Haemophilus influenzae type b vaccine. Int J Infect Dis 7: 143–51

Bagal SK, Brown AD, Cox PJ, Omoto K, Owen RM, Pryde DC, Sidders B, Skerratt SE, Stevens EB, Storer RI, Swain NA (2013) Ion channels as therapeutic targets: a drug discovery perspective. J Med Chem 56: 593–624

Baker PJ, Hiernaux JR, Fauntleroy MB, Stashak PW, Prescott B, Cantrell JL, Rudbach JA (1988) Ability of monophosphoryl lipid A to augment the antibody response of young mice. Infect Immun 56: 3064–6

Bjune G, Høiby EA, Grønnesby JK, Arnesen O, Fredriksen JH, Halstensen A, Holten E, Lindbak AK, Nøkleby H, Rosenqvist E (1991) Effect of outer membrane vesicle vaccine against group B meningococcal disease in Norway. Lancet 338: 1093–6

Blake M, Gotschilch E (1987) Functional and immunologic properties of pathogenic Neisseria surface proteins. In Bacterial outer membranes as model systems, pp 377–400. New York: John Wiley and Sons

Brehony C, Hill DM, Lucidarme J, Borrow R, Maiden MC (2015) Meningococcal vaccine antigen diversity in global databases. Euro Surveill 20

Brehony C, Wilson DJ, Maiden MC (2009) Variation of the factor H-binding protein of Neisseria meningitidis. Microbiology 155: 4155–69

Burton DR (1990) Antibody: the flexible adaptor molecule. Trends Biochem Sci 15: 64–9

Carpenter EP, Beis K, Cameron AD, Iwata S (2008) Overcoming the challenges of membrane protein crystallography. Curr Opin Struct Biol 18: 581–6

Chen VB, Arendall WB, Headd JJ, Keedy DA, Immormino RM, Kapral GJ, Murray LW, Richardson JS, Richardson DC (2010) MolProbity: all-atom structure validation for macromolecular crystallography. Acta Crystallogr D Biol Crystallogr 66: 12–21

Christodoulides M, McGuinness BT, Heckels JE (1993) Immunization with synthetic peptides containing epitopes of the class 1 outer-membrane protein of Neisseria meningitidis: production of bactericidal antibodies on immunization with a cyclic peptide. J Gen Microbiol 139: 1729–38

Christodoulides M, Rattue E, Heckels JE (1999) Effect of adjuvant composition on immune response to a multiple antigen peptide (MAP) containing a protective epitope from Neisseria meningitidis class 1 porin. Vaccine 18: 131–9

Claassen I, Meylis J, van der Ley P, Peeters C, Brons H, Robert J, Borsboom D, van der Ark A, van Straaten I, Roholl P, Kuipers B, Poolman J (1996) Production, characterization and control of a Neisseria meningitidis hexavalent class 1 outer membrane protein containing vesicle vaccine. Vaccine 14: 1001–8

Emsley P, Cowtan K (2004) Coot: model-building tools for molecular graphics. Acta Crystallogr D Biol Crystallogr 60: 2126–32

Exley RM, Shaw J, Mowe E, Sun YH, West NP, Williamson M, Botto M, Smith H, Tang CM (2005) Available carbon source influences the resistance of Neisseria meningitidis against complement. J Exp Med 201: 1637–45.

Finne J, Leinonen M, Mäkelä PH (1983) Antigenic similarities between brain components and bacteria causing meningitis. Implications for vaccine development and pathogenesis. Lancet 2: 355–7

Fletcher LD, Bernfield L, Barniak V, Farley JE, Howell A, Knauf M, Ooi P, Smith RP, Weise P, Wetherell M, Xie X, Zagursky R, Zhang Y, Zlotnick GW (2004) Vaccine potential of the Neisseria meningitidis 2086 lipoprotein. Infect Immun 72: 2088–100

Frank MM, Joiner K, Hammer C (1987) The function of antibody and complement in the lysis of bacteria. Rev Infect Dis 9 Suppl 5: S537–45

Germann T, Bongartz M, Dlugonska H, Hess H, Schmitt E, Kolbe L, Kölsch E, Podlaski FJ, Gately MK, Rüde E (1995) Interleukin-12 profoundly up-regulates the synthesis of antigen-specific complement-fixing IgG2a, IgG2b and IgG3 antibody subclasses in vivo. Eur J Immunol 25: 823–9

Green LR, Eiden J, Hao L, Jones T, Perez J, McNeil LK, Jansen KU, Anderson AS (2016) Approach to the Discovery, Development, and Evaluation of a Novel Neisseria meningitidis Serogroup B Vaccine. Methods Mol Biol 1403: 445–69

Headd JJ, Echols N, Afonine PV, Grosse-Kunstleve RW, Chen VB, Moriarty NW, Richardson DC, Richardson JS, Adams PD (2012) Use of knowledge-based restraints in phenix.refine to improve macromolecular refinement at low resolution. Acta Crystallogr D Biol Crystallogr 68: 381–90

Holst J, Martin D, Arnold R, Huergo CC, Oster P, O’Hallahan J, Rosenqvist E (2009) Properties and clinical performance of vaccines containing outer membrane vesicles from Neisseria meningitidis. Vaccine 27 Suppl 2: B3–12

Hoogerhout P, Donders EM, van Gaans-van den Brink JA, Kuipers B, Brugghe HF, van Unen LM, Timmermans HA, ten Hove GJ, de Jong AP, Peeters CC (1995) Conjugates of synthetic cyclic peptides elicit bactericidal antibodies against a conformational epitope on a class 1 outer membrane protein of Neisseria meningitidis. Infect Immun 63: 3473–8

Hughes-Jones NC, Gardner B (1979) Reaction between the isolated globular sub-units of the complement component C1q and IgG-complexes. Mol Immunol 16: 697–701

Jerse AE, Bash MC, Russell MW (2014) Vaccines against gonorrhea: current status and future challenges. Vaccine 32: 1579–87

Jolley KA, Maiden MC This publication made use of the Neisseria Multi Locus Sequence Typing website (http://pubmlst.org/neisseria/) developed by Keith Jolley and sited at the University of Oxfordl. In BMC Bioinformatics.

Jolley KA, Maiden MC (2010) BIGSdb: Scalable analysis of bacterial genome variation at the population level. BMC Bioinformatics 11: 595

Jongerius I, Lavender H, Tan L, Ruivo N, Exley RM, Caesar JJ, Lea SM, Johnson S, Tang CM (2013) Distinct binding and immunogenic properties of the gonococcal homologue of meningococcal factor h binding protein. PLoS Pathog 9: e1003528

Judd RC (1989) Protein I: structure, function, and genetics. Clin Microbiol Rev 2 Suppl: S41–8

Kaaijk P, van Straaten I, van de Waterbeemd B, Boot EP, Levels LM, van Dijken HH, van den Dobbelsteen GP (2013) Preclinical safety and immunogenicity evaluation of a nonavalent PorA native outer membrane vesicle vaccine against serogroup B meningococcal disease. Vaccine 31: 1065–71

Ke SH, Madison EL (1997) Rapid and efficient site-directed mutagenesis by single-tube ‘megaprimer’ PCR method. Nucleic Acids Res 25: 3371–2

Keller LE, Robinson DA, McDaniel LS (2016) Nonencapsulated Streptococcus pneumoniae: Emergence and Pathogenesis. MBio 7: e01792

Leatherbarrow R, Dwek R (1984) Binding of complement subcomponent Clq to mouse IgG1, IgG2a and IgG2b: A novel Clq binding assay. Molecular Immunology 4: 321–327

Luijkx TA, van Dijken H, Hamstra HJ, Kuipers B, van der Ley P, van Alphen L, van den Dobbelsteen G (2003) Relative immunogenicity of PorA subtypes in a multivalent Neisseria meningitidis vaccine is not dependent on presentation form. Infect Immun 71: 6367–71

Madico G, Welsch JA, Lewis LA, McNaughton A, Perlman DH, Costello CE, Ngampasutadol J, Vogel U, Granoff DM, Ram S (2006) The meningococcal vaccine candidate GNA1870 binds the complement regulatory protein factor H and enhances serum resistance. J Immunol 177: 501–10

Maiden MC (2013) The impact of protein-conjugate polysaccharide vaccines: an endgame for meningitis? Philos Trans R Soc Lond B Biol Sci 368: 20120147

Marrack P, McKee AS, Munks MW (2009) Towards an understanding of the adjuvant action of aluminium. Nat Rev Immunol 9: 287–93

Martin DR, Ruijne N, McCallum L, O’Hallahan J, Oster P (2006) The VR2 epitope on the PorA P1.7-2,4 protein is the major target for the immune response elicited by the strain-specific group B meningococcal vaccine MeNZB. Clin Vaccine Immunol 13: 486–91

Masignani V, Comanducci M, Giuliani MM, Bambini S, Adu-Bobie J, Arico B, Brunelli B, Pieri A, Santini L, Savino S, Serruto D, Litt D, Kroll S, Welsch JA, Granoff DM, Rappuoli R, Pizza M (2003) Vaccination against Neisseria meningitidis using three variants of the lipoprotein GNA1870. J Exp Med 197: 789–99

McGuinness B, Barlow AK, Clarke IN, Farley JE, Anilionis A, Poolman JT, Heckels JE (1990) Deduced amino acid sequences of class 1 protein (PorA) from three strains of Neisseria meningitidis. Synthetic peptides define the epitopes responsible for serosubtype specificity. J Exp Med 171: 1871–82

Michaelsen TE, Ihle Ø, Beckstrøm KJ, Herstad TK, Kolberg J, Høiby EA, Aase A (2003) Construction and functional activities of chimeric mouse-human immunoglobulin G and immunoglobulin M antibodies against the Neisseria meningitidis PorA P1.7 and P1.16 epitopes. Infect Immun 71: 5714–23

Michaelsen TE, Kolberg J, Aase A, Herstad TK, Høiby EA (2004) The four mouse IgG isotypes differ extensively in bactericidal and opsonophagocytic activity when reacting with the P1.16 epitope on the outer membrane PorA protein of Neisseria meningitidis. Scand J Immunol 59: 34–9

Moingeon P, Haensler J, Lindberg A (2001) Towards the rational design of Th1 adjuvants. Vaccine 19: 4363–72

Murphy E, Andrew L, Lee KL, Dilts DA, Nunez L, Fink PS, Ambrose K, Borrow R, Findlow J, Taha MK, Deghmane AE, Kriz P, Musilek M, Kalmusova J, Caugant DA, Alvestad T, Mayer LW, Sacchi CT, Wang X, Martin D et al. (2009) Sequence diversity of the factor H binding protein vaccine candidate in epidemiologically relevant strains of serogroup B Neisseria meningitidis. J Infect Dis 200: 379–89

Murphy TF (2015) Vaccines for Nontypeable Haemophilus influenzae: the Future Is Now. Clin Vaccine Immunol 22: 459–66

Murshudov GN, Vagin AA, Dodson EJ (1997) Refinement of macromolecular structures by the maximum-likelihood method. Acta Crystallogr D Biol Crystallogr 53: 240–55

O’Hallahan J, Lennon D, Oster P (2004) The strategy to control New Zealand’s epidemic of group B meningococcal disease. Pediatr Infect Dis J 23: S293–8

Oomen CJ, Hoogerhout P, Bonvin AM, Kuipers B, Brugghe H, Timmermans H, Haseley SR, van Alphen L, Gros P (2003) Immunogenicity of peptide-vaccine candidates predicted by molecular dynamics simulations. J Mol Biol 328: 1083–9

Oomen CJ, Hoogerhout P, Kuipers B, Vidarsson G, van Alphen L, Gros P (2005) Crystal structure of an Anti-meningococcal subtype P1.4 PorA antibody provides basis for peptide-vaccine design. J Mol Biol 351: 1070–80

Petrovsky N, Aguilar JC (2004) Vaccine adjuvants: current state and future trends. Immunol Cell Biol 82: 488–96

Pizza M, Scarlato V, Masignani V, Giuliani MM, Aricò B, Comanducci M, Jennings GT, Baldi L, Bartolini E, Capecchi B, Galeotti CL, Luzzi E, Manetti R, Marchetti E, Mora M, Nuti S, Ratti G, Santini L, Savino S, Scarselli M et al. (2000) Identification of vaccine candidates against serogroup B meningococcus by whole-genome sequencing. Science 287: 1816–20

Poon PH, Phillips ML, Schumaker VN (1985) Immunoglobulin M possesses two binding sites for complement subcomponent C1q, and soluble 1:1 and 2:1 complexes are formed in solution at reduced ionic strength. J Biol Chem 260: 9357–65

Saukkonen K, Leinonen M, Abdillahi H, Poolman JT (1989) Comparative evaluation of potential components for group B meningococcal vaccine by passive protection in the infant rat and in vitro bactericidal assay. Vaccine 7: 325–8

Schaller A, Troller R, Molina D, Gallati S, Aebi C, Stutzmann Meier P (2006) Rapid typing of Moraxella catarrhalis subpopulations based on outer membrane proteins using mass spectrometry. Proteomics 6: 172–80

Schmitt HJ, von Kries R, Hassenpflug B, Hermann M, Siedler A, Niessing W, Clemens R, Weil J (2001) Haemophilus influenzae type b disease: impact and effectiveness of diphtheria-tetanus toxoids-acellular pertussis (-inactivated poliovirus)/H. influenzae type b combination vaccines. Pediatr Infect Dis J 20: 767–74

Schneider MC, Exley RM, Chan H, Feavers I, Kang YH, Sim RB, Tang CM (2006) Functional significance of factor H binding to Neisseria meningitidis. J Immunol 176: 7566–75

Schneider MC, Prosser BE, Caesar JJ, Kugelberg E, Li S, Zhang Q, Quoraishi S, Lovett JE, Deane JE, Sim RB, Roversi P, Johnson S, Tang CM, Lea SM (2009) Neisseria meningitidis recruits factor H using protein mimicry of host carbohydrates. Nature 458: 890–3

Serruto D, Bottomley MJ, Ram S, Giuliani MM, Rappuoli R (2012) The new multicomponent vaccine against meningococcal serogroup B, 4CMenB: immunological, functional and structural characterization of the antigens. Vaccine 30 Suppl 2: B87–97

Sierra GV, Campa HC, Varcacel NM, Garcia IL, Izquierdo PL, Sotolongo PF, Casanueva GV, Rico CO, Rodriguez CR, Terry MH (1991) Vaccine against group B Neisseria meningitidis: protection trial and mass vaccination results in Cuba. NIPH Ann 14: 195–207; discussion 208-10

Sledge CR, Bing DH (1973) Binding properties of the human complement protein Clq. J Biol Chem 248: 2818–23

Telford JL (2008) Bacterial genome variability and its impact on vaccine design. Cell Host Microbe 3: 408–16

Ulmer JB, Valley U, Rappuoli R (2006) Vaccine manufacturing: challenges and solutions. Nat Biotechnol 24: 1377–83

van den Dobbelsteen G, van Dijken H, Hamstra HJ, Ummels R, van Alphen L, van der Ley P (2004) From HexaMen to NonaMen: expanding a multivalent PorA-based meningococcal outer membrane vesicle vaccine. In International Patholgenic Neisseria Conferrence, p 153. Milwaukee, Wisconsin:

van den Elsen JM, Herron JN, Hoogerhout P, Poolman JT, Boel E, Logtenberg T, Wilting J, Crommelin DJ, Kroon J, Gros P (1997) Bactericidal antibody recognition of a PorA epitope of Neisseria meningitidis: crystal structure of a Fab fragment in complex with a fluorescein-conjugated peptide. Proteins 29: 113–25

van der Ley P, Heckels JE, Virji M, Hoogerhout P, Poolman JT (1991) Topology of outer membrane porins in pathogenic Neisseria spp. Infect Immun 59: 2963–71

van der Ley P, van der Biezen J, Poolman JT (1995) Construction of Neisseria meningitidis strains carrying multiple chromosomal copies of the *porA* gene for use in the production of a multivalent outer membrane vesicle vaccine. Vaccine 13: 401–7

WHO (2014) Antimicrobial resistance: global report on surveillance 2014. In

Wi T, Lahra MM, Ndowa F, Bala M, Dillon JR, Ramon-Pardo P, Eremin SR, Bolan G, Unemo M (2017) Antimicrobial resistance in Neisseria gonorrhoeae: Global surveillance and a call for international collaborative action. PLoS Med 14: e1002344

Winn MD, Ballard CC, Cowtan KD, Dodson EJ, Emsley P, Evans PR, Keegan RM, Krissinel EB, Leslie AG, McCoy A, McNicholas SJ, Murshudov GN, Pannu NS, Potterton EA, Powell HR, Read RJ, Vagin A, Wilson KS (2011) Overview of the CCP4 suite and current developments. Acta Crystallogr D Biol Crystallogr 67: 235–42

Zhu W, Thomas CE, Chen CJ, Van Dam CN, Johnston RE, Davis NL, Sparling PF (2005) Comparison of immune responses to gonococcal PorB delivered as outer membrane vesicles, recombinant protein, or Venezuelan equine encephalitis virus replicon particles. Infect Immun 73: 7558–68

